# A footprint of plant eco-geographic adaptation on the composition of the barley rhizosphere bacterial microbiota

**DOI:** 10.1101/2020.02.11.944140

**Authors:** Rodrigo Alegria Terrazas, Katharin Balbirnie-Cumming, Jenny Morris, Pete E Hedley, Joanne Russell, Eric Paterson, Elizabeth M Baggs, Eyal Fridman, Davide Bulgarelli

**Affiliations:** University of Dundee, Plant Sciences, School of Life Sciences, Dundee, United Kingdom; Cell and Molecular Sciences, The James Hutton Institute, Dundee, United Kingdom; Ecological Sciences, The James Hutton Institute, Aberdeen, United Kingdom; Global Academy of Agriculture and Food Security, University of Edinburgh, Royal (Dick) School of Veterinary Studies, Midlothian, United Kingdom; Institute of Plant Sciences, Agricultural Research Organization (ARO), The Volcani Center, Bet Dagan, Israel

## Abstract

The microbiota thriving in the rhizosphere, the thin layer of soil surrounding plant roots, plays a critical role in plant’s adaptation to the environment. Domestication and breeding selection have progressively differentiated the microbiota of modern crops from the ones of their wild ancestors. However, the impact of eco-geographical constraints faced by domesticated plants and crop wild relatives on recruitment and maintenance of the rhizosphere microbiota remains to be fully elucidated. Here we performed a comparative 16S rRNA gene survey of the rhizosphere of 4 domesticated and 20 wild barley (*Hordeum vulgare*) genotypes grown in an agricultural soil under controlled environmental conditions. We demonstrated the enrichment of individual bacteria mirrored the distinct eco-geographical constraints faced by their host plants. Unexpectedly, Elite varieties exerted a stronger genotype effect on the rhizosphere microbiota when compared with wild barley genotypes adapted to desert environments with a preferential enrichment for members of Actinobacteria. Finally, in wild barley genotypes, we discovered a limited, but significant, correlation between microbiota diversity and host genomic diversity. Our results revealed a footprint of the host’s adaptation to the environment on the assembly of the bacteria thriving at the root-soil interface. In the tested conditions, this recruitment cue layered atop of the distinct evolutionary trajectories of wild and domesticated plants and, at least in part, is encoded by the barley genome. This knowledge will be critical to design experimental approaches aimed at elucidating the recruitment cues of the barley microbiota across a range of soil types.

## Introduction

By 2050 the world’s population is expected to reach 9.5 billion and, to ensure global food security, crop production has to increase by 60% in the same timeframe^1^. This target represents an unprecedented challenge for agriculture as it has to be achieved while progressively decoupling yields from non-renewable inputs in the environment^2^ and amidst climatic modifications which are expected to intensify yield-limiting events, such as water scarcity and drought^3^.

A promising strategy proposes to achieve this task by capitalising on the microbiota inhabiting the rhizosphere, the thin layer of soil surrounding plant roots^4^. The rhizosphere microbiota plays a crucial role in plant’s adaptation to the environment by facilitating, for example, plant mineral uptake^5^ and enhancing plant’s tolerance to both abiotic and biotic stresses^6^.

Plant domestication and breeding selection, which have progressively differentiated modern cultivated crops from their wild relatives^7^, have impacted on the composition and functions of the rhizosphere microbiota^8^. These processes were accompanied by an erosion of the host genetic diversity^9^ and there are growing concerns that, in turn, these limited the metabolic diversity of the microbiota of cultivated plants^10^. Thus, to fully unlock the potential of rhizosphere microbes for sustainable crop production, it is necessary to study the microbiota thriving at the root-soil interface in the light of the evolutionary trajectories of its host plants^11^.

Barley (*Hordeum vulgare* L.), a global crop^12^ and a genetically tractable organism^13^, represents an ideal model to study host-microbiota interactions within a plant domestication framework, due to the fact that wild relatives (*H. vulgare* ssp. *spontaneum*) of domesticated varieties (*H. vulgare* ssp. *vulgare*) are accessible for experimentation^14^. We previously demonstrated that domesticated and wild barley genotypes host contrasting bacterial communities^15^ whose metabolic potential modulates the turn-over of the organic matter in the rhizosphere^16^. However, the impact of eco-geographical constraints faced by domesticated plants and crop wild relatives on recruitment and maintenance of the rhizosphere microbiota remains to be fully elucidated. Tackling this knowledge gap is a key pre-requisite to capitalise on plant-microbiota interactions to achieve the objectives of climate-smart agriculture, in particular sustainably enhancing crop production^17^.

Here we investigated whether exposure to different environmental conditions during evolution left a footprint on the barley’s capacity of shaping the rhizosphere bacterial microbiota. We characterised twenty wild barley genotypes from the ‘B1K’ collection sampled in the Southern levant geographic region, one of the centres of domestication of barley^18,19^. This material represents the three-major barley ‘Ecotypes’ adapted to different habitats in the region^20^: the Golan Heights and northern Galilee, (‘North Ecotype’); the coastal Mediterranean strip, (‘Coast Ecotype’); and the arid regions along the river Jordan and southern Negev (‘Desert Ecotype’). We further subdivided these ‘Ecotypes’ into 5 groups of sampling locations according to the average rainfall of the areas, as a proxy for plant’s adaptation to limiting growth conditions: ‘Coast 1’, ‘Coast 2’, ‘Desert 1’ and ‘Desert 2’ and ‘North’, respectively. (Table 1; Figure 1). These wild barley genotypes were grown in a previously characterised soil, representative of barley agricultural areas of Scotland, under controlled environmental conditions, alongside four cultivated ‘Elite’ varieties encompassing the main usage and genetic diversity of the cultivated germplasm (Table 1). We used an Illumina MiSeq 16S rRNA gene amplicon survey to characterise the microbiota inhabiting the rhizosphere and unplanted soil samples. By using ecological indexes, multivariate statistical analyses and barley genome information we elucidated the impact of eco-geographical constraints and host genetics on the composition of the microbial communities thriving at the barley root-soil interface.

**Table 1.**
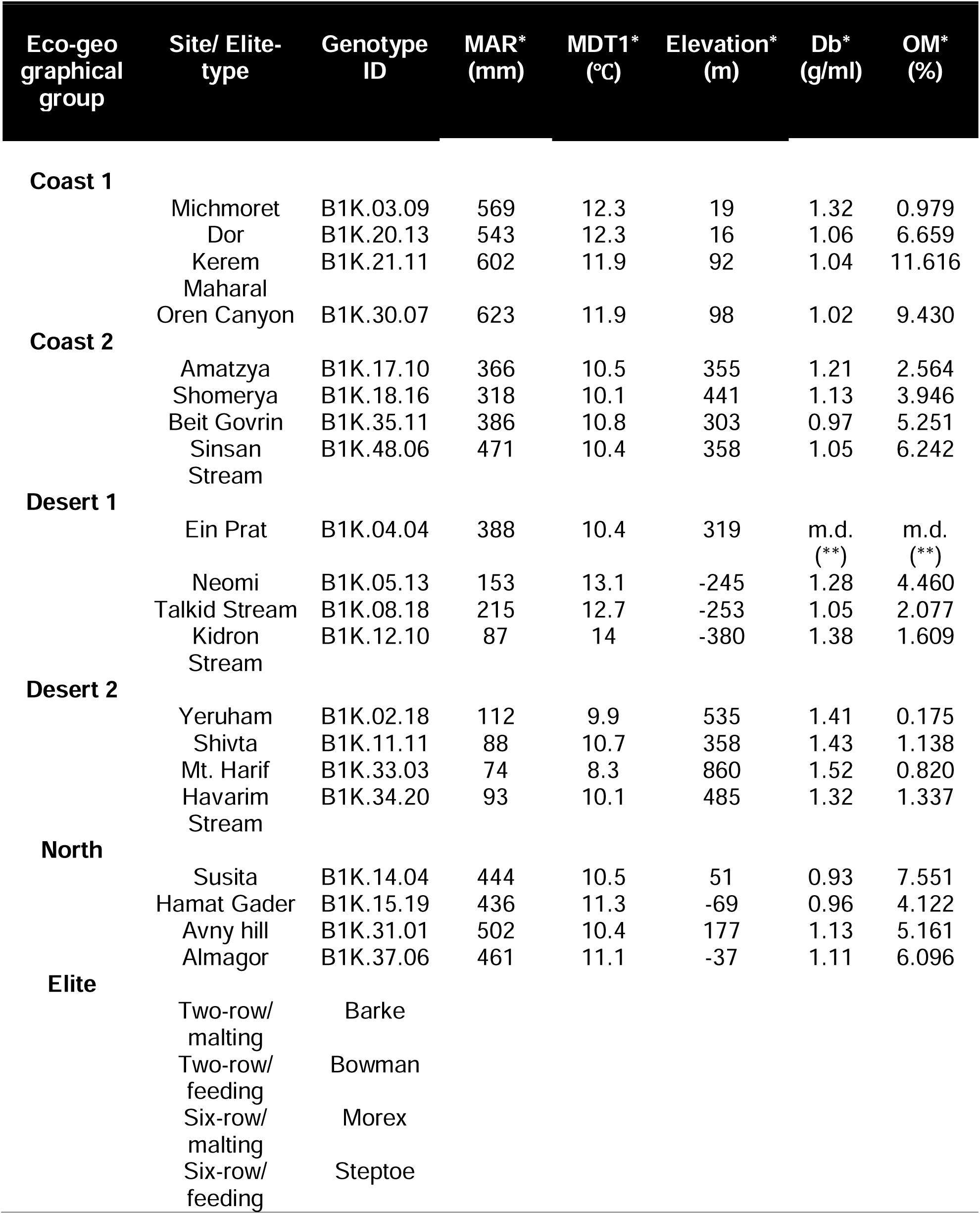
Description of the genotypes used in this study. Eco-geographical group; sampling site or type of the Elite material, genotype ID; mean annual rainfall (MAR*), mid-day temperature in January (MDT1*), Elevation, and soil bulk density (Db*), organic matter content (OM*) of the ‘B1K’ sampling sites from^19,40^. (**) missing data

**Figure 1.**
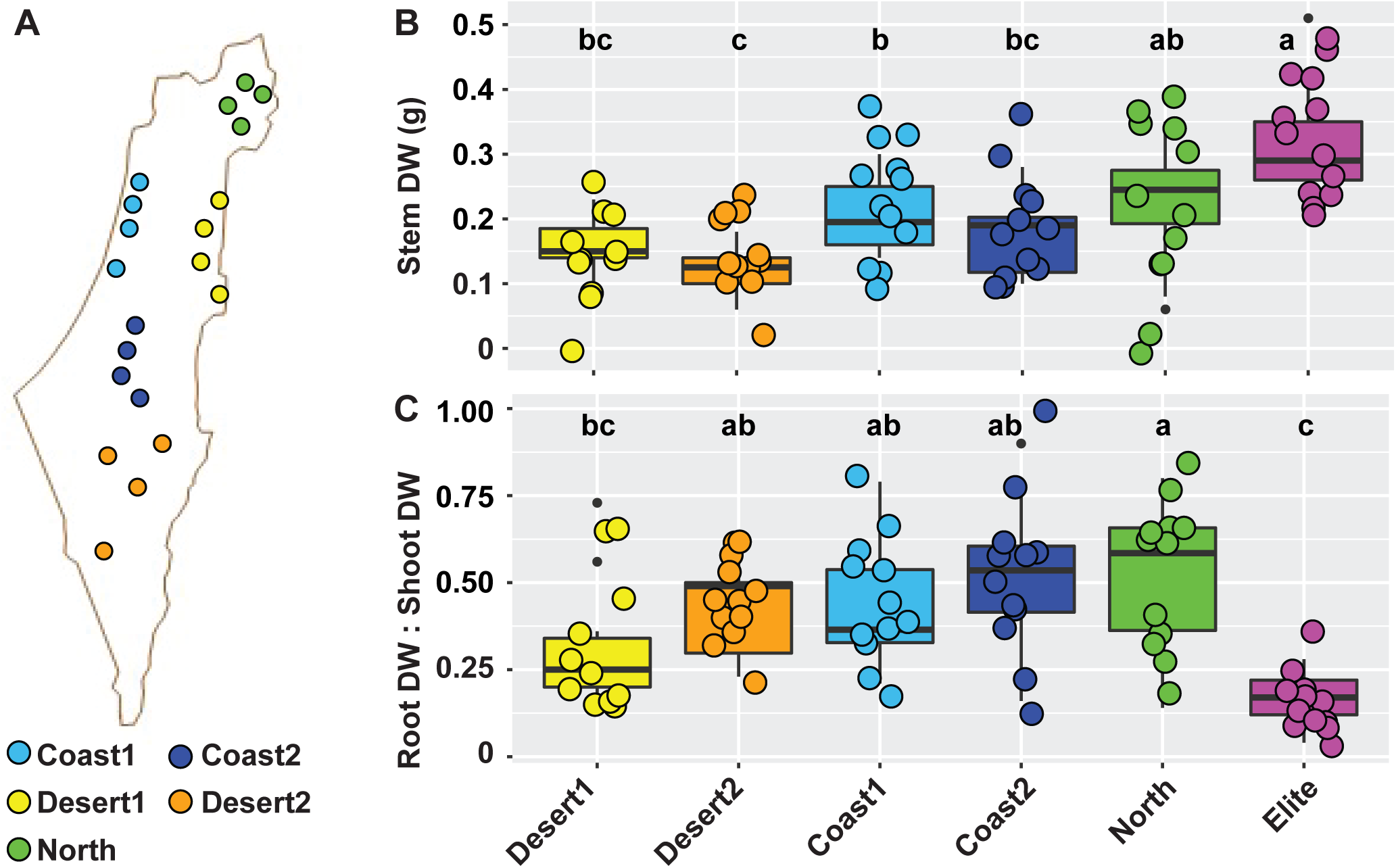
Plant growth parameters of the wild and domesticated barley genotypes. (**a**) Distribution of the twenty wild barley genotypes used in this study in the Israeli geographic region. Individual dots depict the approximate sampling location for a given genotype, colour-coded according to the designated ecogeographic area. (**b**) Stem dry weight of the barley genotypes at the time of sampling. (**c**) Ratio between root and shoot dry weight of the indicated samples. In **b** and **c**, upper and lower edges of the box plots represent the upper and lower quartiles, respectively. The bold line within the box denotes the median, individual shapes depict measurements of individual biological replicates/genotypes for a given group. Different letters within the individual plots denote statistically significant differences between means by Kruskal-Wallis non-parametric analysis of variance followed by Dunn’s post-hoc test (*P* < 0.05).

## Results

### Evolutionary trajectories and eco-geographic adaptation impact on plant growth

Aboveground dry weight from the barley genotypes was measured at early stem elongation as a proxy for plant growth: this allowed us to identify a ‘biomass gradient’ across the tested material. The ‘Elite’ varieties, outperforming wild barley plants, and wild barley genotypes adapted to the more extreme desert environments (i.e., ‘Desert 2’) defined the uppermost and lowermost ranks of this gradient, respectively (*P* < 0.05, Kruskal–Wallis non-parametric analysis of variance followed by Dunn’s post hoc test; Figure 1). Conversely, when we inspected the ratio between above-and belowground biomass we noticed an opposite trend: almost invariably wild barley genotypes allocated more resources than ‘Elite’ varieties to root growth compared to stem growth (*P* < 0.05, Kruskal–Wallis non-parametric analysis of variance followed by Dunn’s post hoc test; Figure 1). As we sampled plants at a comparable developmental stage (Methods; Figure S1), these observations indicate different growth responses of in wild and domesticated genotypes in the tested conditions.

### Taxonomic diversification of the barley microbiota across barley genotypes

To study the impact of these differential responses on the composition of the barley microbiota we generated 6,646,864 16S rRNA gene sequencing reads from 76 rhizosphere and unplanted soil specimens. These high-quality sequencing reads yielded 11,212 Operational Taxonomic Units (OTUs) at 97% identity (Supplementary Dataset 1: worksheet 2). A closer inspection of the taxonomic affiliation of the retrieved OTUs revealed that members of five bacterial phyla, namely *Acidobacteria, Actinobacteria, Bacteroidetes, Proteobacteria* and *Verrucomicrobia*, accounted for more than 97.8% of the observed reads (Figure 2, Supplementary Dataset 1: worksheet 3). Among these dominant phyla, *Bacteroidetes* and *Proteobacteria* were significantly enriched in rhizosphere compared to bulk soil profiles (ANCOM, cut-off 0.6, alpha 0.05, taxa-based corrected, Supplementary Dataset 1: worksheets 4-5).

**Figure 2.**
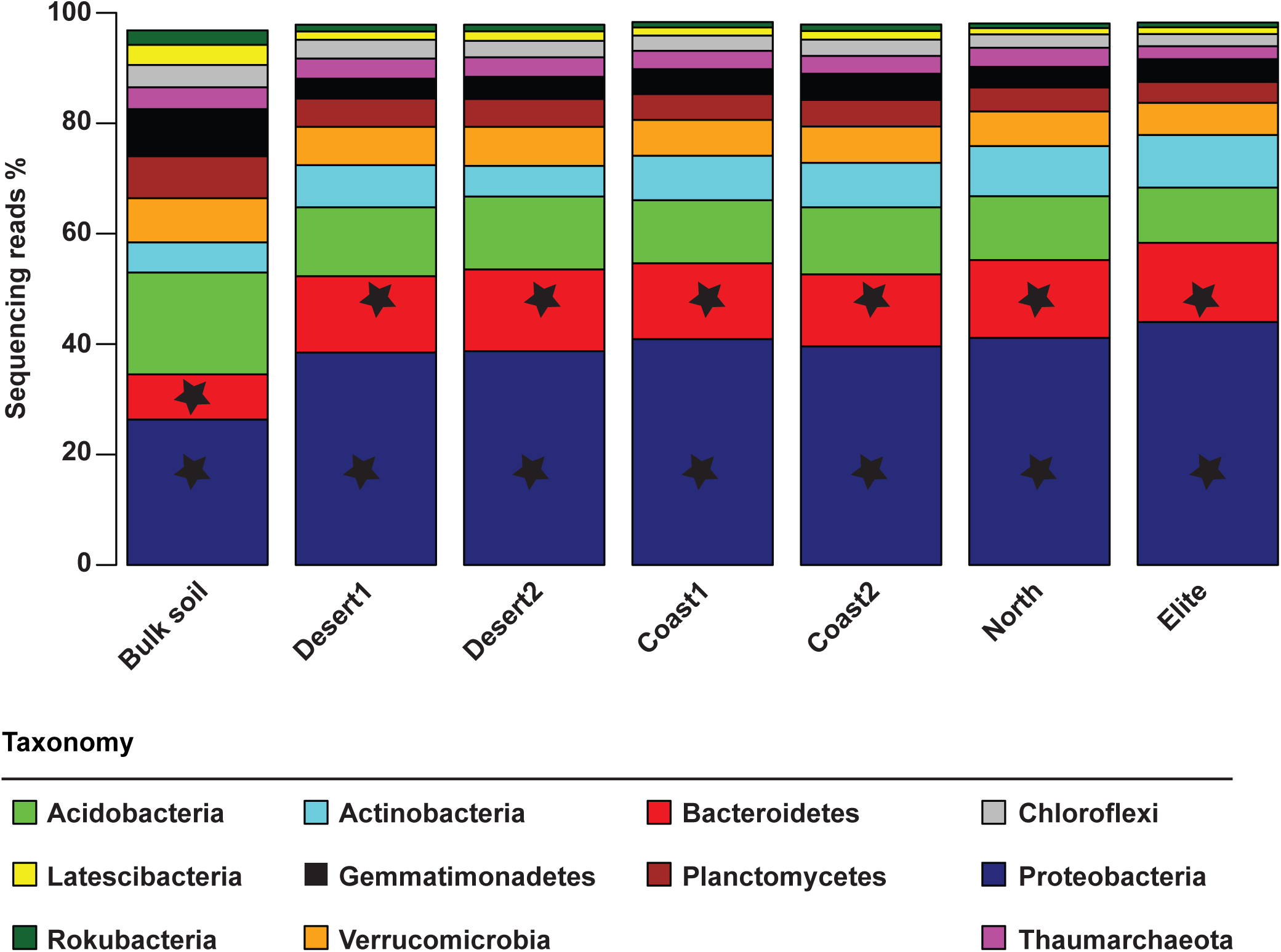
The dominant phyla of the bulk soil and rhizosphere microbiota are conserved across barley genotypes. Average relative abundance (% of sequencing reads) of the dominant phyla retrieved from the microbial profiles of indicated samples. Only phyla displaying a minimum average relative abundance of 1% included in the graphical representation. Stars depict phyla enriched in and discriminating between rhizosphere and between bulk soil samples (ANCOM, cut-off 0.6, alpha 0.05, taxa-corrected).

Next, we investigated the lower ranks of the taxonomic assignments (i.e., OTU level) and computed the Observed OTUs, Chao1 and Shannon indexes for each sample type. This analysis further supported the notion of the rhizosphere as a ‘reduced complexity community’, as both the Observed OTUs and Shannon indexes, but not the projected Chao1, identified significantly richer and more even communities in the bulk soil samples compared to plant-associated specimens (*P* < 0.05, Mann–Whitney U test; Figure S3). Interestingly, when we compared the Chao1 index within rhizosphere samples, we observed that members of the ‘Desert 1’ group assembled a richer and more even community compared with the other genotypes (*P* < 0.05, Kruskal–Wallis non-parametric analysis of variance followed by Dunn’s post hoc test; Figure S3).

To gain further insights into the impact of the sample type on the barley microbiota we generated a canonical analysis of principal coordinates (CAP) using the weighted Unifrac distance, which is sensitive to OTU relative abundance and phylogenetic relatedness. This analysis revealed a marked effect of the microhabitat, i.e., either bulk soil or rhizosphere, on the composition of the microbiota as evidenced by the spatial separation on the axis accounting for the major variation (Figure 3). Interestingly, we observed a clustering of bacterial community composition within rhizosphere samples, which was more marked between ‘Desert’ and ‘Elite’ samples (Figure 3). These observations were corroborated by a permutational analysis of variance which attributed ∼30% of the observed variation to the microhabitat and, within rhizosphere samples, ∼17% of the variation to the individual eco-geographic groups (Permanova *P* <0.01, 5,000 permutations, Table 2). Strikingly similar results were obtained when we computed a Bray-Curtis dissimilarity matrix, which is sensitive to OTUs relative abundance only (Table 2; Figure S4).

**Table 2.** Proportion of rhizosphere microbiota variance explained by the indicated variables and corresponding statistical significance. Levels of the factor Microhabitat are either ‘Bulk soil’ or ‘Rhizosphere’. Levels of the factor Eco-geography are the groups ‘Coast 1’; ‘Coast 2’; ‘Desert 1’; ‘Desert 2’; ‘North’; and ‘Elite’, respectively.

| **Weighted Unifrac** |  |  |
| --- | --- | --- |
| Factor | R2 | Pr(>F) |
| Microhabitat | 0.285 | <0.001 |
| Eco-geography* | 0.168 | <0.001 |
| **Bray-Curtis** |  |  |
| Factor | R2 | Pr(>F) |
| Microhabitat | 0.221 | <0.001 |
| Eco-geography* | 0.129 | <0.001 |

(\*) Analysis performed in rhizosphere samples only

**Figure 3.**
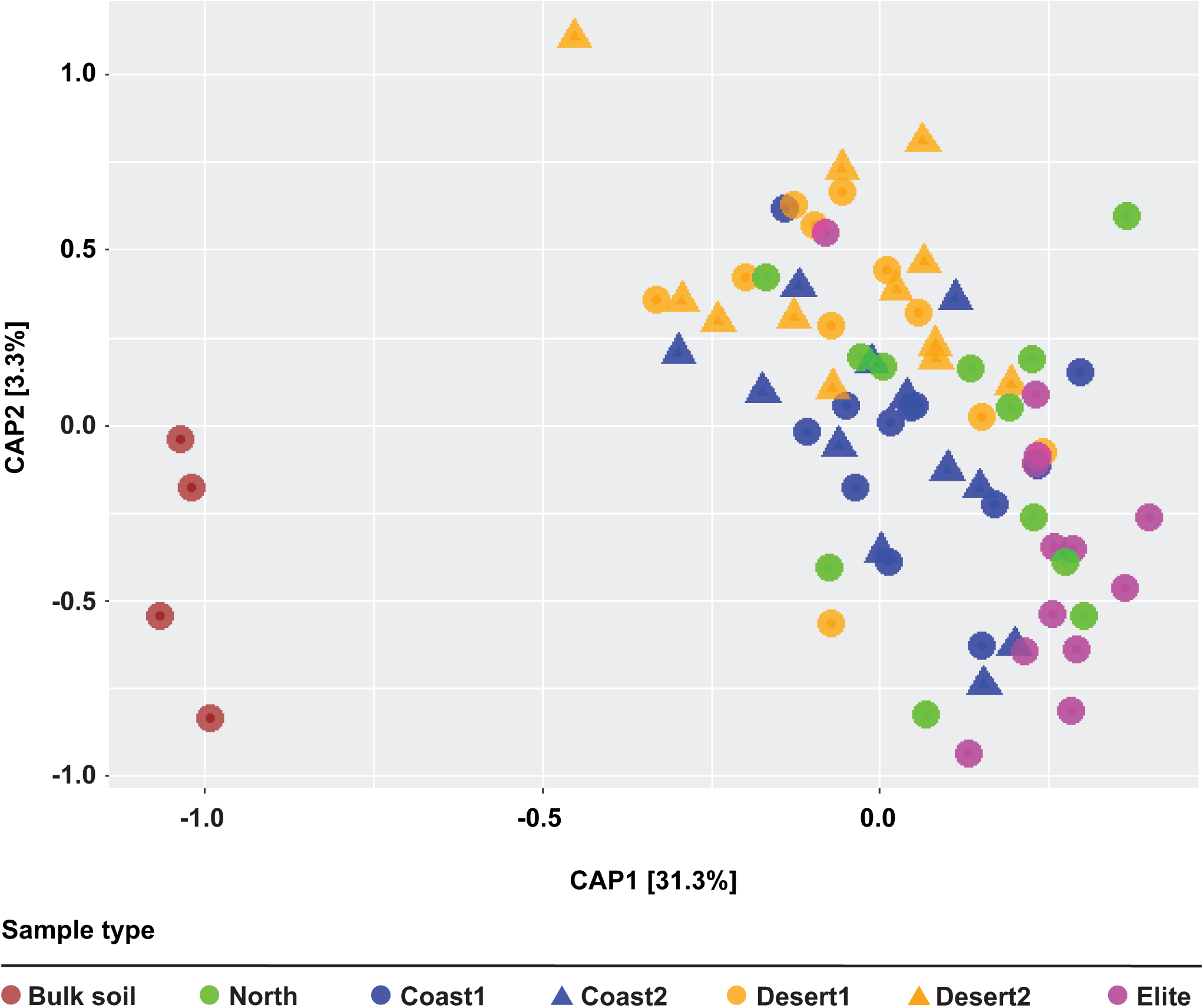
Wild and elite barley genotypes fine-tune the composition of the rhizosphere bacterial microbiota. Principal Coordinates Analysis of the Weighted Unifrac dissimilarity matrix of the microbial communities retrieved from the indicated sample types. Individual shape depicts individual biological replicates colour-coded according to the designated ecogeographic area.

Taken together, these data indicate that the composition of the barley microbiota is fine-tuned by plant recruitment cues which progressively differentiate between unplanted soil and rhizosphere samples and, within these latter, wild ecotypes from elite varieties.

### A footprint of host eco-geographic adaptation shapes the wild barley rhizosphere microbiota

To gain insights into the bacteria underpinning the observed microbiota diversification we performed a series of pair-wise comparisons between ‘Elite’ genotypes and each group of the wild barley ecotypes. This approach revealed a marked specialisation of the members of the ‘Desert’ ecotype compared to ‘Elite’ varieties as evidenced by the number of OTUs differentially recruited between members of these groups (Wald test, *P* <0.05, FDR corrected; Figure 4; Supplementary Dataset 1: worksheets 7-11). Thus, the wild barley ‘Ecotype’ emerged as an element shaping the recruitment cues of the barley rhizosphere microbiota.

**Figure 4.**
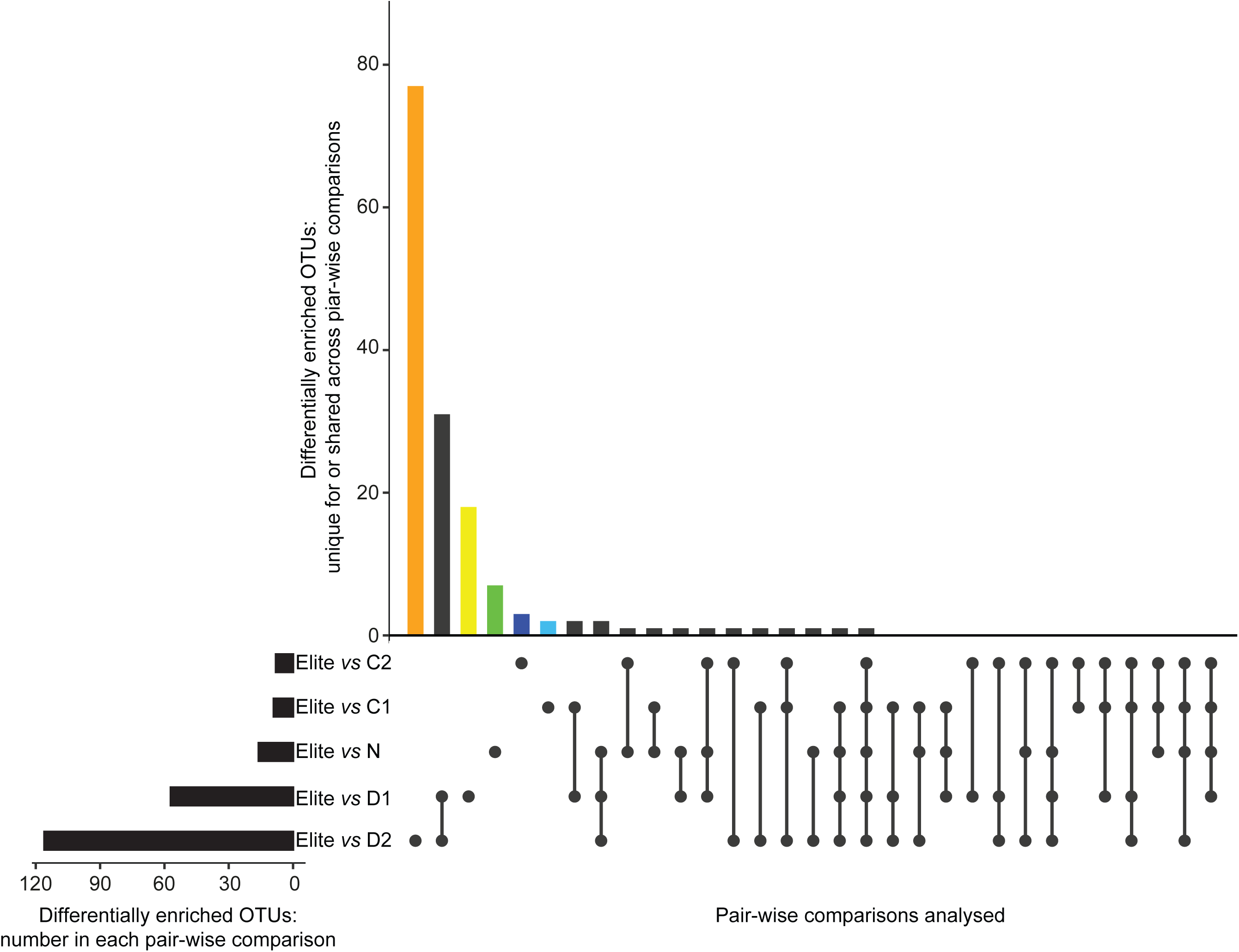
Enrichments of individual bacteria discriminates between elite varieties and wild barley genotypes. Horizontal black bars denote the number of OTUs differentially enriched (Wald test, *P* value < 0.05, FDR corrected) in the indicated pair-wise comparisons between elite varieties and groups of wild barley genotypes. Vertical bars depict the number of differentially enriched OTUs unique for or shared among two or more comparisons highlighted by the interconnected dots underneath the vertical bars. Coloured vertical bars depict differentially enriched OTUs unique for the pair-wise comparisons between ‘Elite’ and ‘Coast 2’ (C2, dark blue), ‘Coast 1’ (C1, light blue), ‘North’ (N, green), ‘Desert 1’ (D1, yellow) and ‘Desert 2’ (D2, orange), respectively.

A closer inspection of the OTUs differentially recruited between ‘Desert’ wild barley and ‘Elite’ varieties revealed that the domesticated material exerted the greatest selective impact on the soil biota, as the majority of the differentially enriched OTUs were enriched in ‘Elite’ varieties (Wald test, *P* <0.05, FDR corrected; Supplementary Dataset 1: worksheets 7 and 8). Next, the taxonomic assignments of these ‘Elite-enriched’ OTUs versus the ‘Desert’ microbiota followed distinct patterns: while the comparison ‘Elite’-‘Desert 1’ produced a subset of enriched OTUs assigned predominantly to *Actinobacteria, Bacteroidetes* and *Proteobacteria*, the comparison ‘Elite’-‘Desert 2’ displayed a marked bias for members of the *Actinobacteria* (i.e., 44 out of 104 enriched OTUs, Figure 5). Consistently, the cumulative abundance of sequencing reads assigned of those Actinobacterial OTUs in ‘Elite’ samples nearly doubled the one recorded for ‘Desert 2’ samples (Figure S5). Within this phylum, we identified a broader taxonomic distribution, as those OTUs were assigned to the families *Intrasporangiaceae, Micrococcaceae, Micromonosporaceae, Nocardioidaceae, Pseudonocardiaceae, Streptomycetaceae*, as well as members of the order *Frankiales*. Interestingly, when we inspect intra-ecotype diversification we identified diagnostic OTUs capable of discriminating between ‘Desert 1’ and ‘Desert 2’ (Wald test, *P* <0.05, FDR corrected; Supplementary Dataset 1: worksheets 12 and 13), while no such a feature was identified discriminating between ‘Coast 1’ and ‘Coast 2’ at the statistical test imposed. Taken together, our data indicate that wild barley ‘Ecotype’ (i.e., the differential effect of ‘North’, ‘Coast, and ‘Desert’ versus ‘Elite’) acts as a determinant for the rhizosphere barley microbiota whose composition is ultimately fine-tuned by a sub-specialisation within the ‘Ecotype’ itself (i.e., the differential effect of ‘Desert 1’ and ‘Desert 2’).

**Figure 5.**
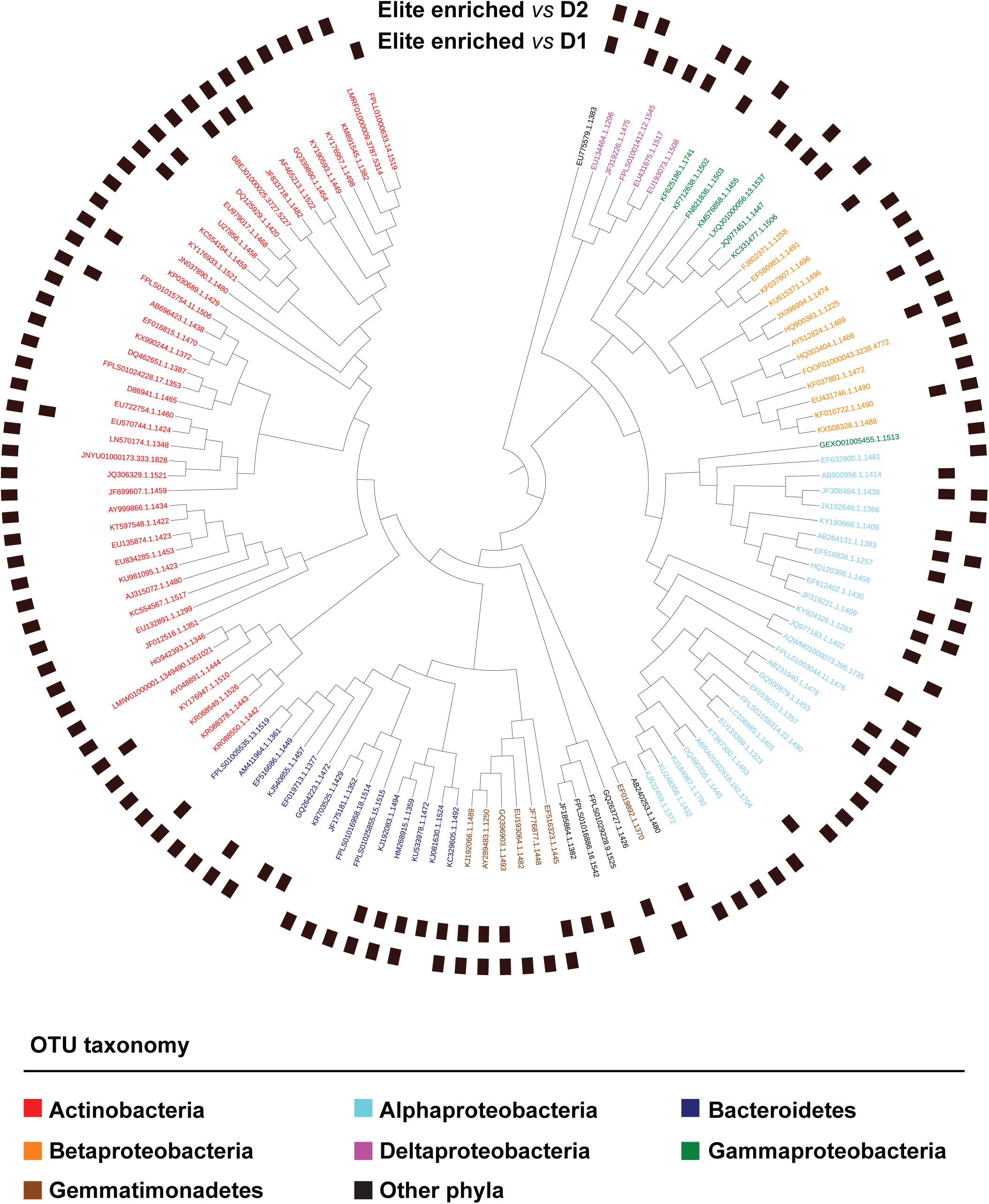
Actinobacteria are preferentially enriched in and discriminate between elite genotypes and wild barley genotypes adapted to desert environments. Individual external nodes of the tree represent one of the OTUs enriched in the rhizosphere of elite genotypes compared to either (or both) rhizosphere samples desert areas (Wald test, *P* value < 0.05, FDR corrected). The colours reflect OTUs’ taxonomic affiliation at Phylum level. A black bar in the outer rings depicts whether that given OTU was identified in the comparisons between ‘Elite’ and either ‘Desert 1’ or ‘Desert 2’ genotypes, respectively. Phylogenetic tree constructed using OTUs 16S rRNA gene representative sequences.

These observations prompted us to investigate whether the differential microbiota recruitment between the tested plants was encoded, at least in part, by the barley genome. We therefore generated a dissimilarity matrix using Single Nucleotide Polymorphisms (SNPs) available for the tested genotypes and we inferred their genetic relatedness using a simple matching coefficient (Supplementary Dataset 1: worksheet 14). With few notable exceptions, this analysis revealed three distinct clusters of genetically related plants, represented by and reflecting the ‘Elite’ material, the ‘Desert’ and the ‘Coast’ wild barley genotypes (Figure S6). The genetic diversity between domesticated material exceeded their microbial diversity (compare relatedness of “Elite” samples in Figure 3 with the ones of Figure S6) as further evidenced by the fact that we failed to identify a significant correlation between these parameters (*P* value > 0.05). However, when we focused the analysis solely on the pool of wild barley genotypes, we obtained a significant correlation between genetic and microbial distances (Mantel test r = 0.230; *P* value < 0.05; Figure 6).

**Figure 6:**
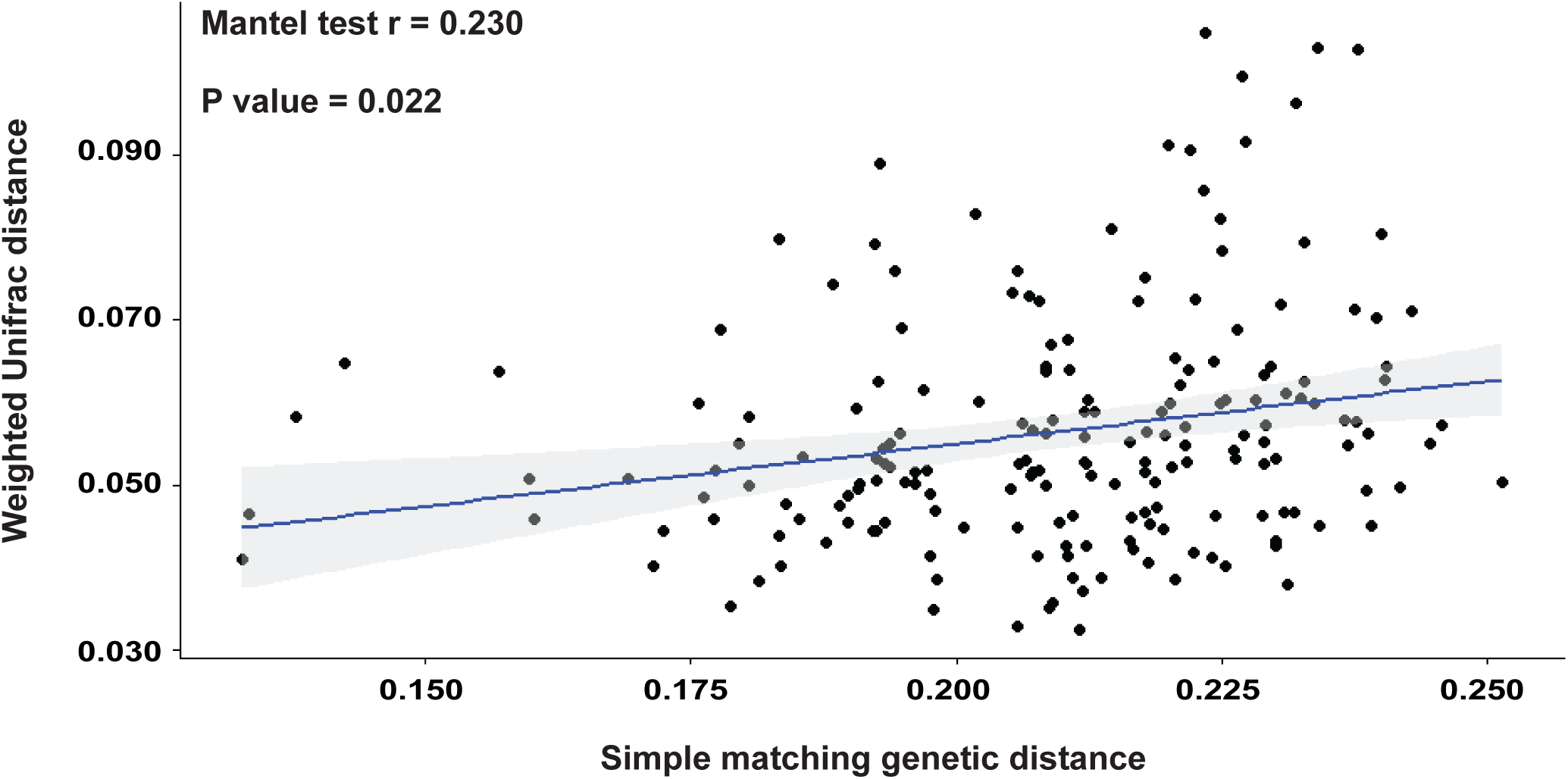
Mantel test between genetic distance and microbial distance in the wild barley rhizosphere. Individual dots depict individual comparison of any given pair of wild barley genotypes between average value of weighted unifrac distance (y-axis) and genetic distance shown as simple matching coefficients (x-axis). The blue line depicts the regression line, the grey shadow the 95% confidence interval, respectively.

Taken together, this revealed a footprint of barley host’s adaptation to the environment on the assembly of the bacteria thriving at the root-soil interface. This recruitment cue interjected the distinct evolutionary trajectories of wild and domesticated plants and, at least in part, is encoded by the barley genome.

## Discussion

In this study we investigated how plant genotypes adapted to different eco-geographic niches may recruit a distinct microbiota once exposed to a common environment.

As we performed a ‘common environment experiment’ in a Scottish agricultural soil, we first determined how the chosen experimental conditions related to the ones witnessed by wild barleys in their natural habitats. Strikingly, the aboveground biomass gradient observed in our study, with ‘Elite’ material almost invariably outperforming wild genotypes and material sampled at the locations designated ‘Desert 2’ at the bottom of the ranking, “matched” the phenotypic characterisation of members of the ‘B1K’ collection grown in a ‘common garden experiment’ in a local Israeli soil^18^. Conversely, belowground resource allocation followed an opposite pattern as evidenced by an increased root:shoot dry weight ratio in wild genotypes compared to ‘Elite’ varieties. As responses to edaphic stress, such as drought tolerance, may modulate the magnitude of above-belowground resource partitioning in plants^21^ and root traits^22^, our data might reflect the adaptation of the wild barley exposure to dry areas. Taken together, these results suggest adaptive that responses to eco-geographic constraints in barley have a genetic inheritance component which can be detected and studied in controlled conditions.

As genetically-inherited root traits have been implicated in shaping the rhizosphere microbiota in barley^23^ and other crops^24^, these observations motivated us to examine whether these below-ground differences were reflected by changes in microbiota recruitment. The distribution of reads assigned to given phyla appears distinct in plant-associated communities which are dominated in terms of abundance by members of the phyla *Acidobacteria, Actinobacteria, Bacteroidetes and Proteobacteria*, with these two latter phyla significantly enriched in rhizosphere samples compared to bulk soil controls. This taxonomic affiliation is consistent with previous investigations in barley in either the same^25^ or in a different soil type^15^ as well as in other crop plants^26^. In summary, these data indicate that the higher taxonomic ranks of the barley rhizosphere microbiota are conserved across soil types as well as wild and domesticated genotypes.

The characterisation of the microbiota at lower taxonomic ranks, i.e., the OTU-level, revealed a significant effect of the microhabitat (i.e., either bulk soil or rhizosphere) and, within plant-associated communities, a footprint of eco-geographic adaptation. For instance, alpha diversity indexes clearly pointed at selective processes modulating bacterial composition as the number of Observed OTUs and the Shannon index indicate simplified and reduced-complexity communities inhabiting the rhizosphere compared to unplanted soil. This can be considered a hallmark of the rhizosphere microbiota as it has been observed in multiple plant species and across soils^6^. Conversely, within rhizosphere samples, alpha-diversity analysis failed to identify a clear pattern, except for the Chao1 index revealing a potential for a richer community associated with plants sampled at the ‘Desert 1’ locations. This motivated us to further explore the between-sample diversity, which is beta-diversity. This analysis revealed a clear host-dependent diversification of the bacteria associated to barley plants manifested by ∼17% of the variance of the rhizosphere microbiota explained by the eco-geographical location of the sampled material. This value exceeded the host genotype effect on the rhizosphere microbiota we previously observed in wild and domesticated barley plants^15^, but is aligned with the magnitude of host effect observed in the rhizosphere microbiota of modern and ancestral genotypes of rice^27^ and common bean^28^. As these studies were conducted in different soil types, our data suggest that the magnitude of host control on the rhizosphere microbiota is ultimately fine-tuned by and in response to soil characteristics.

The identification of the bacteria underpinning the observed microbiota diversification led to three striking observations. First, the comparison between ‘Elite’ varieties and the material representing the ‘Desert’ ecotype was associated with the largest number of differentially recruited OTUs, while the other wild barley genotypes appeared to share a significant proportion of their microbiota with domesticated plants. A prediction of this observation is that the distinct evolutionary trajectories of wild and domesticated plants *per se* cannot explain the host-mediated diversification of the barley microbiota. As aridity and temperature played a prominent role in shaping the phenotypic characteristics of barley^19,29^ it is tempting to speculate that the adaptation to these environmental parameters played a predominant role also in shaping microbiota recruitment.

Second, it is the domesticated material which exerted a stronger effect on microbiota recruitment, manifested by the increased number of host-enriched OTUs compared to wild barley genotypes. This suggests that the capacity of shaping the rhizosphere microbiota has not been “lost” during barley domestication and breeding selection. Our findings are consistent with data gathered for domesticated and ancestral common bean genotypes, which revealed that shifts from native soils to agricultural lands led to a stronger host-dependent effect on rhizosphere microbes^30^. Due to the intrinsic limitation of 16S rRNA gene profiles of predicting the functional potential of individual bacteria, it will be necessary to complement this investigation with whole-genome surveys^31,32^ and metabolic analyses^16,33^ to fully discern the impact of the host genotype on the functions provided by the rhizosphere microbiota to their hosts.

The third observation is the marked quantitative enrichment of OTUs assigned to the phylum *Actinobacteria* in ‘Elite’ varieties when compared to members of the ‘Desert’ ecotype, in particular plants of the ‘Desert 2’ locations. At first glance, the ‘direction’ of this bacterial enrichment is difficult to reconcile with the eco-geographic adaptation of wild barleys and, in particular, the fact that *Actinobacteria* are more tolerant to arid conditions^34^ and, consequently, more abundant in desert vs. non-desert soils^35^. However, the enrichment of *Actinobacteria* in modern crops compared to ancestral relatives has recently emerged as a distinctive feature of the microbiota of multiple plant species^36^. Although the ecological significance of this trait of the domesticated microbiota remains to be fully elucidated, studies conducted in rice^37^ and other grasses, including barley^38^, indicate a relationship between drought stress and *Actinobacteria* enrichments. These observations suggest that the wild barley genome has evolved the capacity to recognise microbes specifically adapted to the local conditions and, in turn, to repress the growth of others. For instance, among the bacteria differentially enriched between ‘Desert 1’ and ‘Desert 2’ we identified genera, such as *Arthrobacter sp*., adapted to extreme environments and long-term nutrient starvation^39^, possibly reflecting the differential adaptation of ‘Desert 1’ and ‘Desert 2’ plants to soil with limited organic matter^40^.

Interestingly, we were able to trace the host genotype effect on rhizosphere microbes to the genome of wild barley. This suggests that, similar to other wild species^11^, microbiota recruitment co-evolved with other adaptive traits. Conversely, the genetic diversity in ‘Elite’ material largely exceeded microbiota diversity. This is reminiscent of studies conducted in maize which failed to identify a significant correlation between polymorphisms in the host genome and alpha-and beta-diversity characteristics of the rhizosphere microbiota^41,42^. Yet, and again similar to maize^43^, our data indicate that the recruitment of individual bacterial OTUs in the ‘Elite’ varieties, rather than community composition as a whole, is the feature of the rhizosphere microbiota under host genetic control.

Although these findings were gathered from the individual soil tested and further validation across a range of soil types is required, a prediction from these observations is that the host control of the rhizosphere microbiota is exerted by a limited number of loci in the genome with a relatively large effect. This is congruent with our previous observation that mono-mendelian mutations in a single root trait, root hairs, impact on ∼18% of the barley rhizosphere microbiota^25^.

Likewise, this scenario is compatible with a limited number of genes controlling the biosynthesis and rhizodeposition of defensive secondary metabolites which have been implicated in shaping the plant microbiota^44^. Among these compounds, the indol-alkaloids benzoxazinoids recently gained centre-stage as master regulators of the maize-associated microbial communities^45-47^. Interestingly, *Hordeum vulgare* has evolved a distinct indol-alkaloid compound, gramine^48^, which is preferentially accumulated in the tissues of the wild genotypes compared to ‘Elite’ varieites^49^ and whose physiological properties are comparable to the ones of benzoxazinoids^50^. Whether gramine or other species-specific secondary metabolites contribute, at least in part, to shape the barley microbiota will be the focus of future investigations.

Since modern varieties have been selected with limited or no knowledge of belowground interactions, how was the capacity of shaping the rhizosphere microbiota retained within the cultivated germplasm? The recent observation that genes controlling reproductive traits display pleiotropic effects on root system architecture^51^ could provide a direct link between crop selection and microbiota recruitment in modern varieties. These traits, and in particular genes encoding flower developments, show a marked footprint of eco-geographic adaptation and have been selected during plant domestication and breeding^29^. By manipulating those genes, breeders may have manipulated also belowground traits, and in turn, the microbiota thriving at the root-soil interface. With an increased availability of genetic^52^ and genomic^53^ resources for wild and domesticated barleys, this hypothesis can now be experimentally tested and the adaptive significance of the barley rhizosphere microbiota ultimately deciphered. Specifically, interspecific populations within the wild^54^ as well as between wild and cultivated^52^ germplasm, could be deployed in genetic mapping experiments aimed at identifying barley genetic determinants of the rhizosphere microbiota.

## Conclusions

Our results revealed a footprint of host’s adaptation to the environment on the assembly of the bacteria thriving at the root-soil interface in barley. This recruitment cue layered atop of the distinct evolutionary trajectories of wild and domesticated plants and, at least in part, is encoded by the barley genome. Although our study was limited to the individual soil investigated, our sequencing survey will provide a reference dataset for the development of indexed bacterial collections of the barley microbiota. These can be used to infer causal relationships between microbiota composition and plant traits, as demonstrated for *Arabidopsis thaliana*^55^ and rice^56^. Furthermore, this knowledge is critical for the establishment of reciprocal transplantation experiments aimed at elucidating the adaptive value of crop-microbiota interactions, similar to what has recently been proposed for the model plant *A. thaliana*^57^. However, for crop plants like barley, this will necessarily be conditioned by two elements: identifying the host genetic determinants of the rhizosphere microbiota and inferring microbial metabolic potential *in situ*. Ultimately, this will help devising strategies aimed at sustainably enhancing crop production for climate-smart agriculture.

## Supporting information

Supplementary Dateset 1

## Methods

### Soil

The soil was sampled from the agricultural research fields of the James Hutton Institute, Invergowrie, Scotland, UK in the Quarryfield site (56°27’5"N 3°4’29"W; Sandy Silt Loam, pH 6.2; Organic Matter 5%; Table S1). This field was left unplanted and unfertilised in the three years preceding the investigations and previously used for barley-microbiota interactions investigations^25^.

### Plant genotypes

Twenty wild barley genotypes (*Hordeum vulgare* ssp. *spontaneum*) and four ‘Elite’ cultivars (*Hordeum vulgare* ssp. *vulgare*) were used and described in Table 1. Wild barley genotypes were selected representing eco-geographical variation of the ‘B1K’ collection^18,19^. The ‘Elite’ genotypes were selected as a representation of different types of spring barley in plant genetic studies. The cultivar ‘Morex’ is an American six-row malting variety whose genome was the first to be sequenced^58^. The cultivars ‘Bowman’ and ‘Barke’ are two-row varieties, developed in US for feed and in Germany for malting, respectively, whereas Steptoe is an American six-row type used for animal feed^52,59,60^.

### Plant growth conditions

Barley seeds were surface sterilized as previously reported^61^ and germinated on 0.5% agar plates at room temperature. Seedlings displaying comparable rootlet development after 5 days post-plating were sown individually in 12-cm diameter pots containing approximately 500g of the ‘Quarryfield’ soil, plus unplanted pots filled with bulk soil as controls. Plants were arranged in a randomised design with this number of replicates: ‘Coast1’ number of replicates *n*=12; ‘Coast2’ *n*=12; ‘Desert1’ *n*=11; ‘Desert2’ *n*=12; ‘North’ *n*=12; ‘Elite’ *n*=13 (Supplementary Dataset 1: worksheet 1). Plants were grown for 5 weeks in a glasshouse at 18/14 °C (day/night) temperature regime with 16 h day length and watered every two days with 50 ml of deionized water.

### Bulk soil and rhizosphere DNA preparation

At early stem elongation, corresponding to Zadoks stages 30-32 ^62^, plants were pulled from the soil and the stems and leaves were separated from the roots (Figure S1). Above-ground plant parts were dried at 70 °C for 72 h and the dry weight recorded. The roots were shaken manually to remove excess of loosely attached soil. For each barley plant, the top 6 cm of the seminal root system and the attached soil layer was collected and placed in sterile 50 ml falcon tube containing 15 ml phosphate-buffered saline solution (PBS). Rhizosphere was operationally defined, for these experiments, as the soil attached to this part of the roots and extracted through this procedure. The samples were then vortexed for 30s and aseptically transferred to a second 50ml falcon containing 15ml PBS and vortexed again for 30s to ensure the dislodging and suspension of the rhizosphere soil. Then, the two falcon tubes with the rhizosphere suspension were mixed and centrifuged at 1,500 x *g* for 20min, the supernatant was removed, with the rhizosphere soil collected as the pellet, flash frozen with liquid nitrogen and stored at −80°C, until further use. After the rhizosphere extraction step, these parts of the roots were combined with the rest of the root system for each plant, thoroughly washed with water removing any attached soil particles and dried at 70°C for 72h for root biomass measurement. Bulk soil samples were collected from the 6cm below the surface of unplanted pots and subjected to the same procedure as above.

DNA was extracted from the rhizosphere samples using FastDNA SPIN Kit for Soil (MP Biomedicals, Solon, USA) according to the manufacturer’s recommendations. The concentration and quality of DNA was checked using a Nanodrop 2000 (Thermo Fisher Scientific, Waltham, USA) spectrophotometer and stored at −20°C until further use. DNA concentration was used as a proxy for the proportion of the sampled microbiota and evaluated across sample type (Figure S2).

### Preparation of 16 rRNA gene amplicon pools

The hypervariable V4 region of the small subunit rRNA gene was the target of amplification using the PCR primer pair 515F (5’-GTGCCAGCMGCCGCGGTAA-3’) and 806R (5’-GGACTACHVGGGTWTCTAAT-3’). The PCR primers had incorporated an Illumina flow cell adapter at their 5’ end and the reverse primers contained 12bp unique ‘barcode’ for simultaneous sequencing of several samples^63^. PCR, including No-Template Controls (NTCs) for each barcoded primer, was performed as previously reported with the exception of the BSA at 10mg/ml concentration per reaction^25^. Only samples whose NTCs yielded an undetectable PCR amplification were retained for further analysis.

### Illumina 16S rRNA gene amplicon sequencing

The pooled amplicon library was submitted to the Genome Technology group, The James Hutton Institute (Invergowrie, UK) for quality control, processing and sequencing as previously described^25,64-66^. Briefly, samples were sequenced using an Illumina MiSeq platform with the 2 × 150bp chemistry.

### Sequencing reads processing

Sequencing reads were processed and analysed using a custom bioinformatics pipeline. First, QIIME (Quantitative Insights into Microbial Ecology) software, version 1.9.0, was used to process the FASTQ files following default parameters for each step^67^. The forward and reverse read files from individual libraries were decompressed and merged using the command join_paired_ends.py, with a minimum overlap of 30bp between reads. Then, the reads were demultiplexed according to the barcode sequences. Quality filtering was performed using the command split_libraries_fastq.py, imposing a minimum acceptable PHRED score ‘-q’ of 20. Next, these high quality reads were truncated at the 250^th^ nucleotide using the function ‘fastq_filter’ implemented in USEARCH^68^. Only these high-quality PE, length-truncated reads were used for clustering in Operational Taxonomic Units (OTUs) a 97% sequence identity. OTUs were identified using the ‘closed reference’ approach against Silva database (version 132)^69^. OTU-picking against the Silva database was performed using the SortMeRNA algorithm^70^, producing in an OTU table containing the abundance of OTUs per sample plus a phylogenetic tree. To control for potential contaminant OTUs amplified during library preparation, we retrieved a list of potential environmental contaminant OTUs previously identified in our laboratory^65^ and we used this list to filter the results of the aforementioned OTU-enrichment analysis. Additionally, singleton OTUs, (OTUs accounting for only one sequencing read in the whole dataset) and OTUs assigned to chloroplast and mitochondria (taken as plant derived sequences) were removed using the command filter_otus_from_otu_tables.py. Taxonomy matrices, reporting the number of reads assigned to individual phyla, were generated using the command summarize_taxa.py. The OTU table, the phylogenetic tree and the taxonomy matrix, were further used in R for visualizations and statistical analysis.

### Statistical analyses I: univariate datasets and 16S rRNA gene alpha and beta-diversity calculations

Analysis of the data was performed in R^71^ using a custom script with the following packages: Phyloseq^72^ for processing, Alpha and Beta-diversity metrics; ggplot2 ^73^ for data visualisations; Vegan^74^ for statistical analysis of beta-diversity; Ape^75^ for phylogenetic tree analysis. For any univariate dataset used (e.g., aboveground biomass; DNA concentration) the normality of the data’s distribution was checked using Shapiro–Wilk test. Non-parametric analysis of variance were performed by Kruskal-Wallis Rank Sum Test, followed by Dunn’s post hoc test with the functions kruskal.test and the posthoc.kruskal.dunn.test, respectively, from the package PMCMR.

For Alpha-diversity analysis, the OTU table was rarefied at 11,180 reads per sample and this allowed us to retain 8,744 OTUs for downstream analyses (Supplementary Dataset 1: worksheet 6). The Chao1, Observed OTUs and Shannon indices calculated using the function estimate richness in Phyloseq package. Beta-diversity was analysed using a normalized OTU table (i.e., not rarefied) for comparison. For the construction of the normalized OTU table, low abundance OTUs were further filtered removing those not present at least 5 times in 20% of the samples, to improve reproducibility. Then, to control for the uneven number of reads per specimen, individual OTU counts in each sample were divided over the total number of generated reads for that samples and converted in counts per million. Beta-diversity was analysed using two metrics: Bray-Curtis that considers OTUs relative abundance and Weighted Unifrac that additionally is sensitive to phylogenetic classification^76^. These dissimilarity matrices were visualized using Canonical Analysis of Principal coordinates (CAP)^77^ using the ordinate function in the Phyloseq package and its significance was inspected using a permutational ANOVA over 5,000 permutations.

Beta-diversity dissimilarity matrices were assessed by Permutational Multivariate Analysis of Variance (Permanova) using Adonis function in Vegan package over 5,000 permutations to calculate effect size and statistical significance.

### Statistical analyses II: analysis of Phyla and OTUs differentially enriched among samples

The analysis of the Phyla whose abundances differentiated among rhizosphere and bulk soil samples was performed with analysis of composition of microbiomes (ANCOM)^78^ imposing 0.6 cut-off and 0.05 alpha value (taxa-based corrected) as previously described^79^.

The analysis of the OTUs whose abundances differentiated among samples was performed a) between individual eco-geographic groups and bulk soil samples to assess the rhizosphere effect and b) between the rhizosphere samples to assess the eco-geographic effect. The eco-geographic effect was further corrected for a microhabitat effect (i.e., for each group, only OTUs enriched against both unplanted soil and at least another barley genotype were retained for further analysis). The analysis was performed using the DESeq2 method^80^ with an adjusted *P* value < 0.05 (False Discovery Rate, FDR corrected). This method was selected since it outperforms other hypothesis-testing approaches when data are not normally distributed and a limited number of individual replicates per condition (i.e., approximately 10) are available^81^. DESeq2 was performed using the eponymous named package in R with the OTU table filtered for low abundance OTUs as an input.

The number of OTUs differentially recruited in the pair-wise comparisons between ‘Elite’ and wild barley genotypes was visualised using the package UpSetR^82^.

The phylogenetic tree was constructed using the representative sequences of the OTUs significantly differentiating ‘Elite’ genotypes and either ‘Desert1’ or ‘Desert2’ samples annotated with iTOL^83^.

### Statistical analyses III: Correlation plot genetic distance-microbial distance

To assess the genetic variation on the barley germplasm we used the SNP platform ‘BOPA1’^84^ comprising 1,536 single nucleotide polymorphisms. We used GenAlex 6.5^85,86^ to construct a genetic distance matrix using the simple matching coefficient. Genetic distance for the barley genotypes was visualised by hierarchical clustering using the function hclust in R. Microbial distance was calculated on the average distances for each ecogeographic group using the Weighted Unifrac metric. Correlation between the plant’s genetic and microbial distances was performed using a mantel test with the mantel.rtest of the package ade4 in R. The correlation was visualised using the functions ggscatter of the R packages ggpbur.

## Availability of Materials and Data

The sequences generated in the 16S rRNA gene sequencing survey are deposited in the European Nucleotide Archive (ENA) under the accession number PRJEB35359. The version of the individual packages and scripts used to analyse the data and generate the figures of this study are available at https://github.com/BulgarelliD-Lab/Barley_B1K

## Acknowledgements

We are grateful to Prof Andy Flavell (University of Dundee) for providing us with the ‘B1K’ seeds used in this study. We thank Malcolm Macaulay for the technical assistance during the sequencing library preparation. We thank Dr Timothy George (The James Hutton Institute) for the critical comments on the manuscript.

## Authors information

### Contributions

The study was conceived by RAT and DB with critical inputs from EP and EB. RAT and KBC performed the experiments. JM and PH generated the 16S rRNA sequencing reads. JR provided access to the molecular marker information of the barley genome. EF provided access to the eco-geographical and phenotypic data of the B1K accessions. RAT and DB analysed the data. All authors critically reviewed and edited the manuscript and approved its publication.

### Corresponding author

Correspondence to Davide Bulgarelli

