## Supplementary Information for "A footprint of plant eco-geographic adaptation on the composition of the barley rhizosphere bacterial microbiota"

Rodrigo Alegria Terrazas^1^, Katharin Balbirnie-Cumming^1^, Jenny Morris^2^, Pete E Hedley^2^, Joanne Russell^2^, Eric Paterson^3^, Elizabeth M Baggs^4^, Eyal Fridman^5^ and Davide Bulgarelli^1*^

^1^University of Dundee, Plant Sciences, School of Life Sciences, Dundee, United Kingdom;

^2^Cell and Molecular Sciences, The James Hutton Institute, Dundee, United Kingdom;

^3^Ecological Sciences, The James Hutton Institute, Aberdeen, United Kingdom;

^4^Global Academy of Agriculture and Food Security, University of Edinburgh, Royal (Dick) School of Veterinary Studies, Midlothian, United Kingdom.

^5^Institute of Plant Sciences, Agricultural Research Organization (ARO), The Volcani Center, Bet Dagan, Israel.

***Correspondence:**
Davide Bulgarelli

### **Description of Supplementary Dataset 1 content**

**Worksheet 1**: Experimental design and sample description for the 16S rRNA gene sequencing experiment. **Worksheet 2**: OTUs and individual counts identified the amplicon sequencing survey. **Worksheet 3**: Matrix depicting the phylum relative abundances of the microbes identified in the amplicon sequencing survey. **Worksheet 4**: Individual cumulative read abundances at phylum level for ANCOM analysis. **Worksheet 5**: ANCOM output. **Worksheet 6**: OTUs and individual counts of the rarefied table. **Worksheets 7-11**: Taxonomic information of the OTUs differentially recruited in the pairwise comparison ‘Elite’ and ‘Desert 1’; ‘Elite’ and ‘Desert 2’; ‘Elite’ and ‘Coast 1’; ‘Elite’ and ‘Coast 2’; ‘Elite’ and ‘North’, respectively. **Worksheets 12-13**: Taxonomic information of the OTUs differentially enriched in ‘Desert 1’ versus ‘Desert 2’; and ‘Desert 2’ versus ‘Desert 1’, respectively. **Worksheet 14**: Genetic information of the tested germplasm. **Worksheet 15**: Mapping file of the samples for the genetic distance/microbial distance analysis. **Worksheet 16**: Mapping file of the samples for the ANCOM analysis.

Table S1: Chemical and physical characteristic of the ‘Quarryfield’ soil used in this study.

| Organic matter (%) | 5.0 |
| --- | --- |
| Soil particles (%) |  |
| Silt | 47.39 |
| Clay | 11.28 |
| Sand | 41.33 |
| Soil texture | Sandy Silt Loam |
| pH^1^ | 6.2 |
| Mineral content (ppm) |  |
| Phosphorus^2^ | 99 |
| Potassium | 162 |
| Magnesium | 156 |
| Calcium | 2,661 |
| Nitrogen content (mg Kg^-1^) |  |
| Ammonium | 1.8 |
| Nitrate | 13.5 |

1-Determined using water as extractant. 2-Determined using the Olsen method.

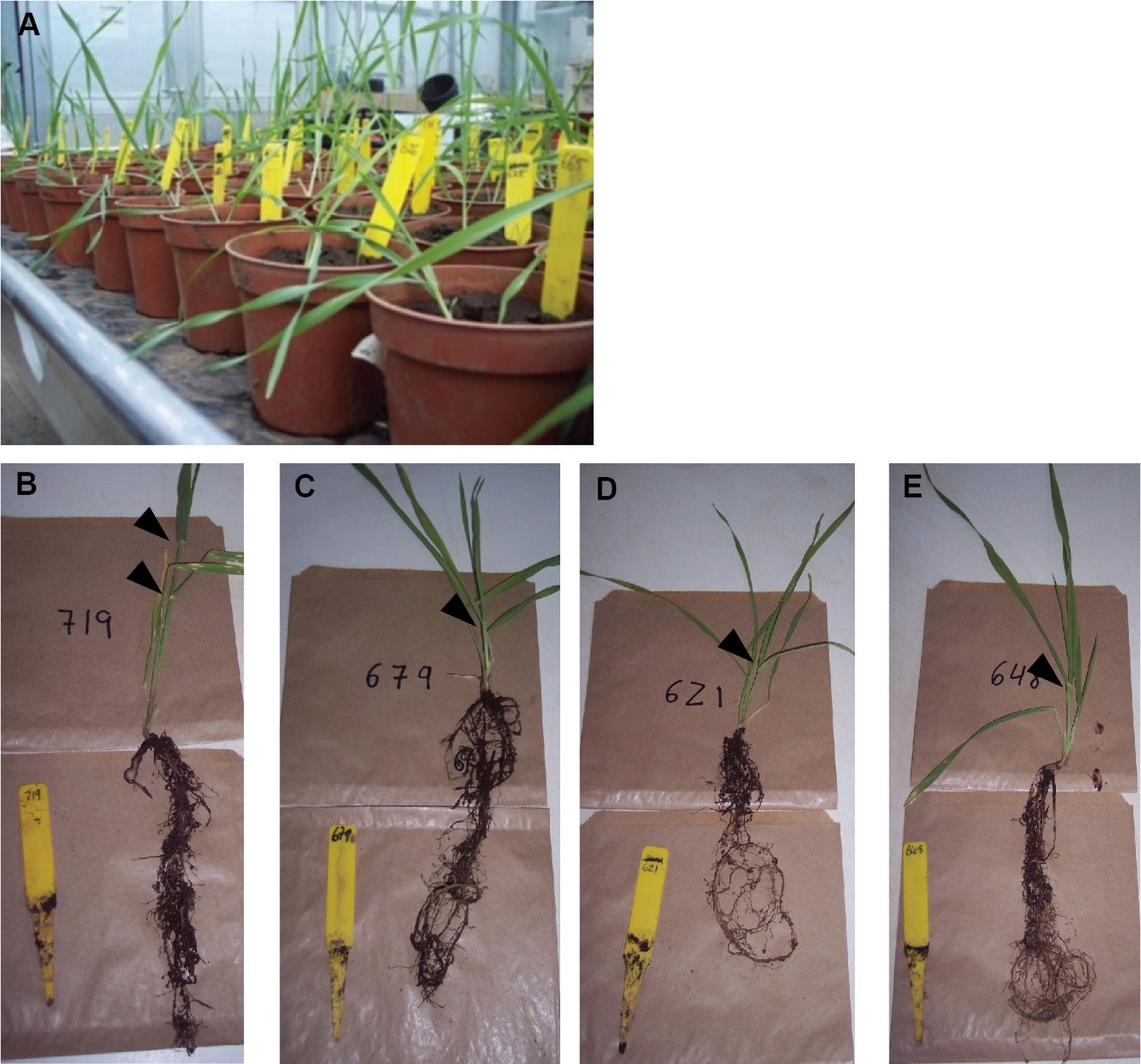

Figure S1: Above- and Below-ground developmental characteristics of the barley plants used in this study. (A) Barley pots during the glasshouse growth. Individual sampled plants from (B) ‘Elite’, (C) ‘Coast’, (D) ‘Desert’ and (E) ‘North’ displaying the whole root system prior fractionation of the uppermost 6cm. Arrowheads indicate detectable nodes per stem. In panels B-E, label width = 1.5 cm; Label length = 13 cm.

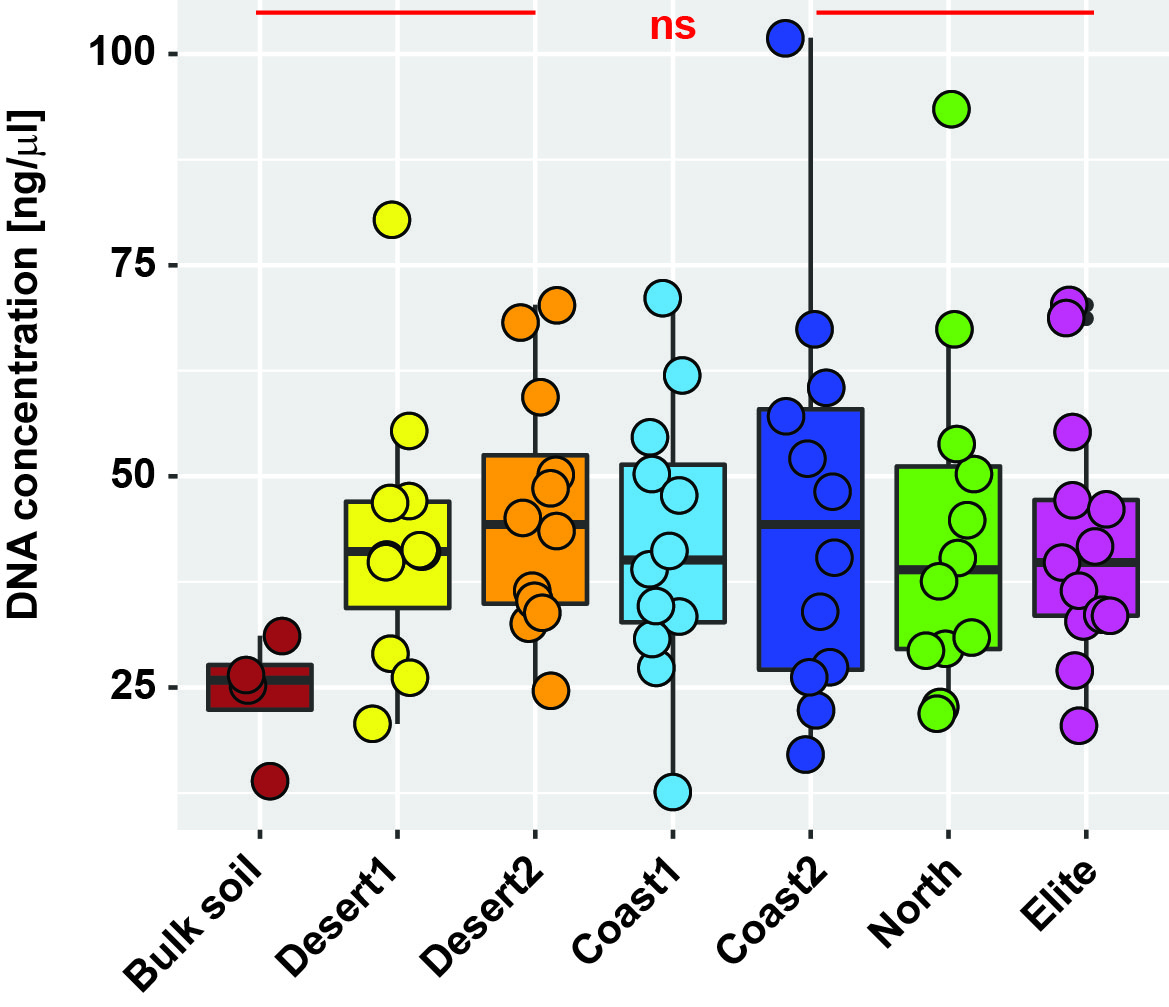
**Figure S2: DNA concentration recorded from the bulk soil and rhizosphere samples.** Individual dots depict individual biological samples for the indicated group; ns, no significant differences at *P* < 0.05 observed among means by Kruskal-Wallis non-parametric analysis of variance followed by Dunn’s post-hoc test.

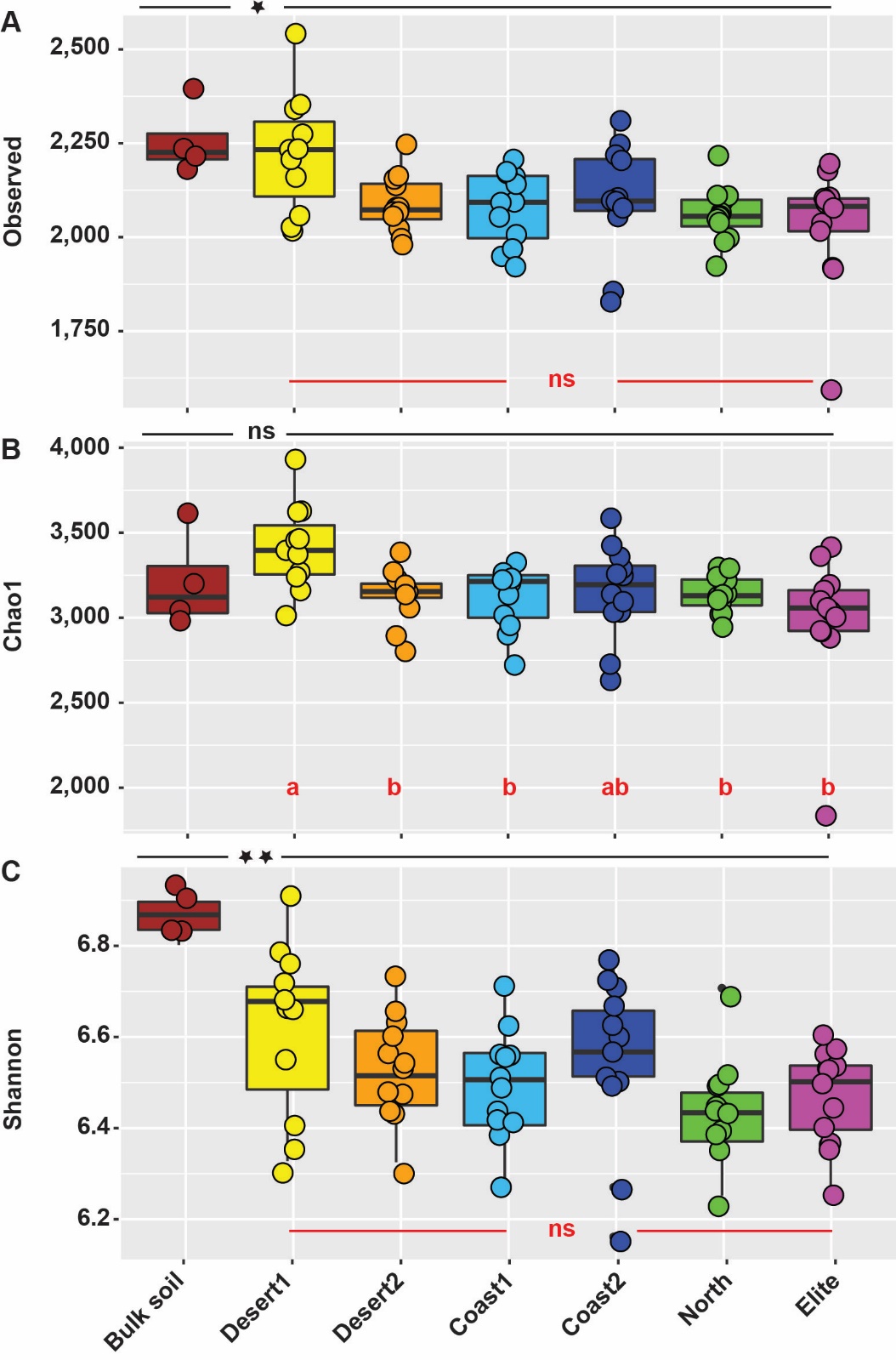

Figure S3: Alpha-diversity calculations of Observed OTUs (A), Chao 1 (B), and Shannon (C) recorded for the indicated eco-geographic groups and ‘Elite’ genotypes Data computed using OTUs clustered at 97% similarity. Asterisks denote statistically significant differences between microhabitat by non-parametric Wilcoxon rank sum test (* *P* < 0.05; ** *P* < 0.01; ns not significant). Different red letters within rhizosphere samples denote statistically significant differences between barley groups means by Kruskal-Wallis non-parametric analysis of variance followed by Dunn’s post-hoc test (*P* < 0.05); ns, no significant differences observed at *P* < 0.05.

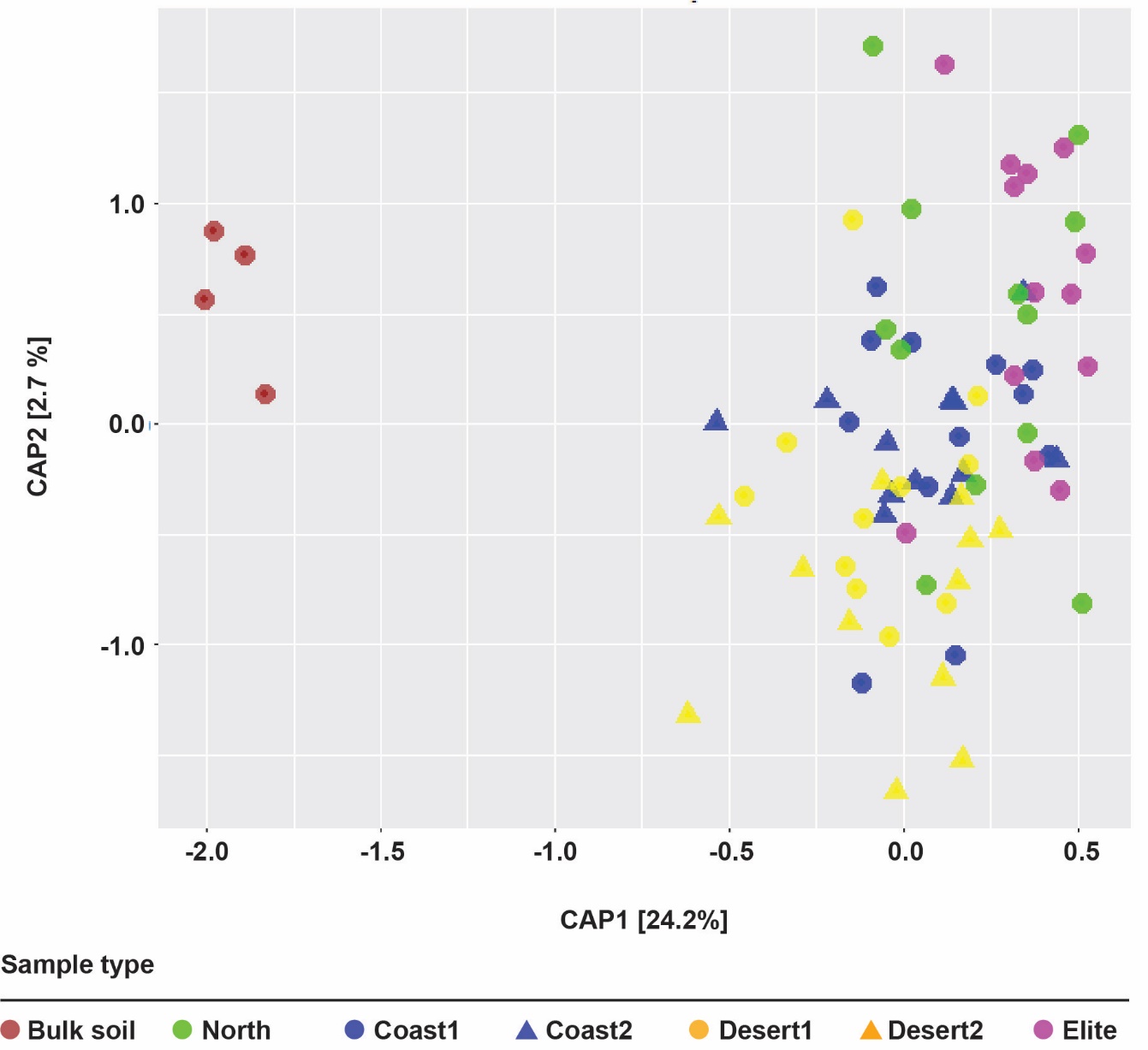

**Figure S4: Wild and ‘Elite’ barley genotypes fine-tune the composition of the rhizosphere bacterial microbiota**. Principal Coordinates Analysis of the Bray-Curtis dissimilarity matrix of the microbial communities retrieved from the indicated sample types. Individual shape depicts individual biological replicates colour-coded according to the designated eco-geographic area.

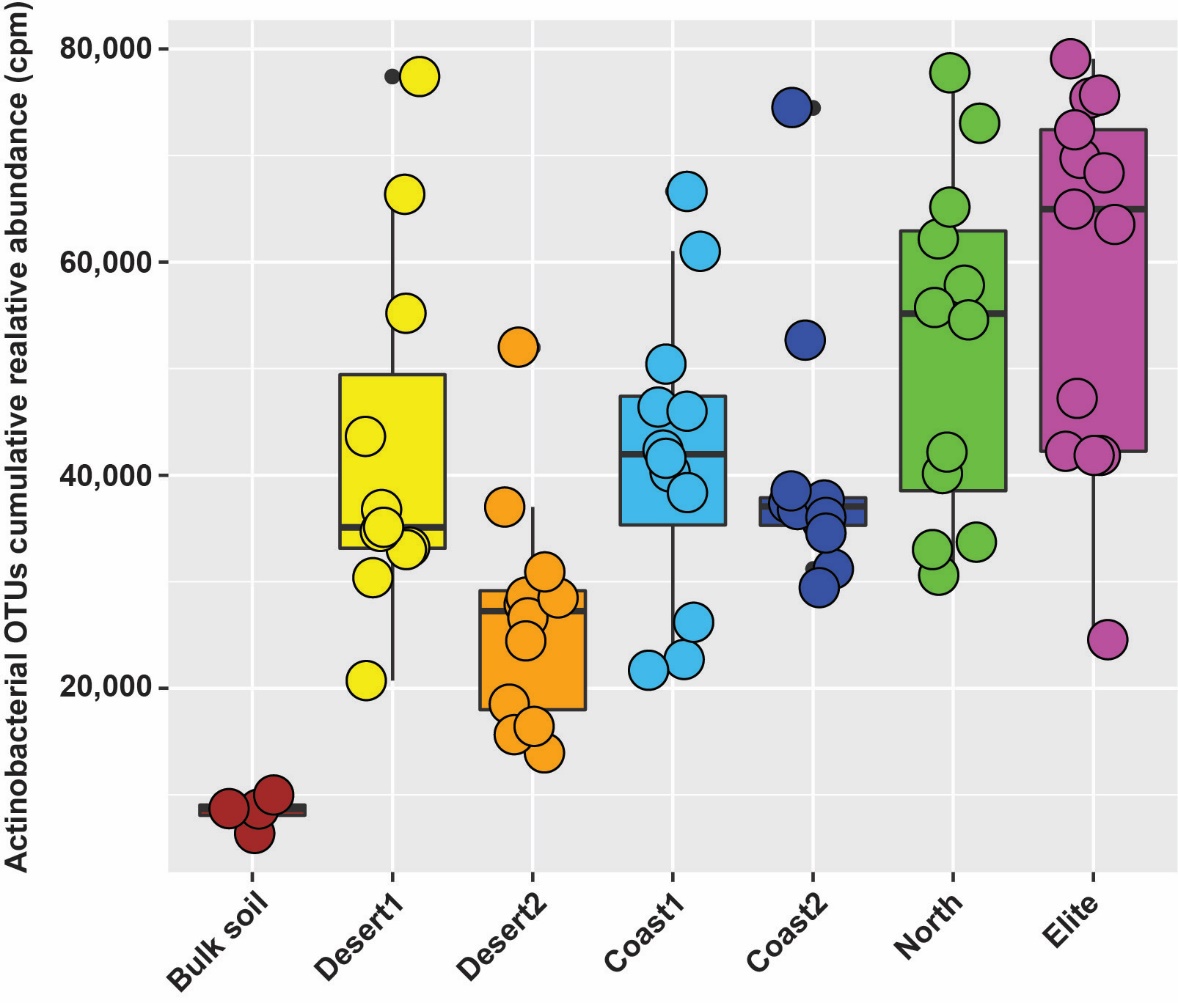

**Figure S5: Distribution of Elite-enriched Actinobacterial OTUs across sample type**. Cumulative relative abundances of the sequencing reads assigned to OTUs classified as ‘Actinobacteria’ and significantly enriched in ‘Elite’ versus wild barley genotypes (Wald test, P <0.05, FDR corrected). Individual dots depict individual replicates assigned to a given sample type.

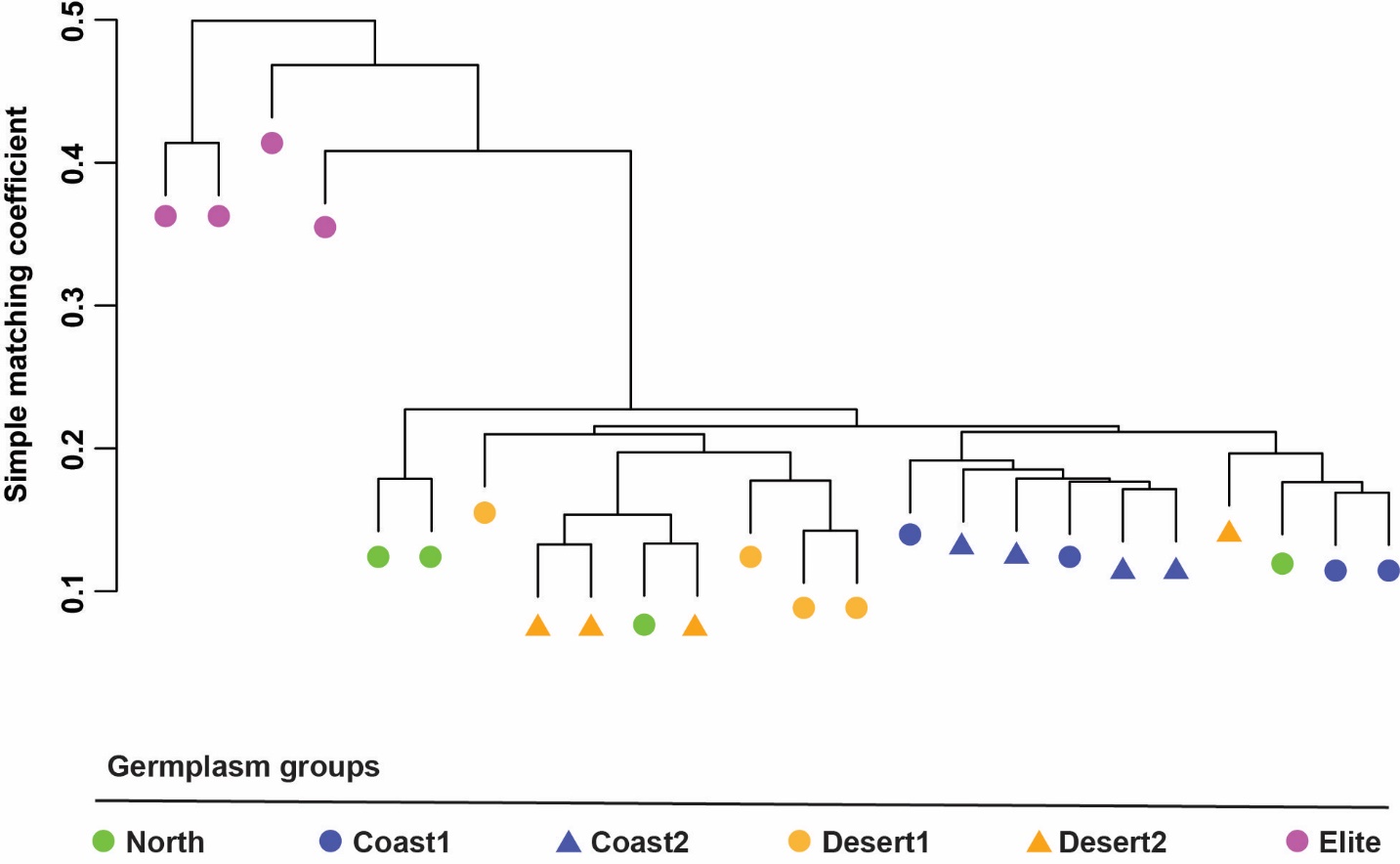

**Figure S6: Genetic relatedness of the individual barley genotypes used in this study.** Hierarchical clustering of the genetic distance expressed as simple matching coefficient of the indicated barley samples colour-coded according their ecogeographic area. Genetic distance computed using the SNP information of the BOPA1 markers.
